# Iterative Hard Thresholding in GWAS: Generalized Linear Models, Prior Weights, and Double Sparsity

**DOI:** 10.1101/697755

**Authors:** Benjamin B. Chu, Kevin L. Keys, Christopher A. German, Hua Zhou, Jin J. Zhou, Eric Sobel, Janet S. Sinsheimer, Kenneth Lange

## Abstract

1

**Background:** Consecutive testing of single nucleotide polymorphisms (SNPs) is usually employed to identify genetic variants associated with complex traits. Ideally one should model all covariates in unison, but most existing analysis methods for genome-wide association studies (GWAS) perform only univariate regression.

**Results:** We extend and efficiently implement iterative hard thresholding (IHT) for multiple regression, treating all SNPs simultaneously. Our extensions accommodate generalized linear models (GLMs), prior information on genetic variants, and grouping of variants. In our simulations, IHT recovers up to 30% more true predictors than SNP-by-SNP association testing, and exhibits a 2 to 3 orders of magnitude decrease in false positive rates compared to lasso regression. We also test IHT on the UK Biobank hypertension phenotypes and the Northern Finland Birth Cohort of 1966 cardiovascular phenotypes. We find that IHT scales to the large datasets of contemporary human genetics and recovers the plausible genetic variants identified by previous studies.

**Conclusions:** Our real data analysis and simulation studies suggest that IHT can (a) recover highly correlated predictors, (b) avoid over-fitting, (c) deliver better true positive and false positive rates than either marginal testing or lasso regression, (d) recover unbiased regression coefficients, (e) exploit prior information and group-sparsity and (f) be used with biobank sized data sets. Although these advances are studied for GWAS inference, our extensions are pertinent to other regression problems with large numbers of predictors.

## 2 Introduction

In genome-wide association studies (GWAS), modern genotyping technology coupled with imputation algorithms can produce an *n* × *p* genotype matrix **X** with *n* ≈ 10^6^ subjects and *p* ≈ 10^7^ genetic predictors (10; 41). Data sets of this size require hundreds of gigabytes of disk space to store in compressed form. Decompressing data to floating point numbers for statistical analyses leads to matrices too large to fit into standard computer memory. The computational burden of dealing with massive GWAS datasets limits statistical analysis and interpretation. This paper discusses and extends a class of algorithms capable of meeting the challenge of multiple regression models with modern GWAS data scales.

Traditionally, GWAS analysis has focused on SNP-by-SNP (single nucleotide polymorphism) association testing (10; 9), with a p-value computed for each SNP via linear regression. This approach enjoys the advantages of simplicity, interpretability, and a low computational complexity of 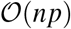. Furthermore, marginal linear regressions make efficient use of computer memory, since computations are carried out on genotype *vectors* one at a time, as opposed to running on the full genotype *matrix* in multiple regression. Some authors further increase association power by reframing GWAS as a linear mixed model problem and proceeding with variance component selection (20; 26). These advances remain within the scope of marginal analysis.

Despite their numerous successes (41), marginal regression is less than ideal for GWAS. It implicitly assumes that all SNPs have independent effects. In contrast, multiple regression can in principle model the effect of all SNPs simultaneously. This approach captures the biology behind GWAS more realistically because traits are usually determined by multiple SNPs acting in unison. Marginal regression selects associated SNPs one by one based on a pre-set threshold. Given the stringency of the p-value threshold, marginal regression can miss many causal SNPs with low effect sizes. As a result, heritability is underestimated. When *p* ≫ *n*, one usually assumes that the number of variants *k* associated with a complex trait is much less than n. If this is true, we can expect multiple regression models to perform better because it a) offers better outlier detection (35) and better prediction, b) accounts for the correlations among SNPs, and c) allows investigators to model interactions. Of course, these advantages are predicated on finding the truly associated SNPs.

Adding penalties to the loss function is one way of achieving parsimony in multiple regression. The lasso (39; 40) is the most popular model selection device in current use. The lasso model selects non-zero parameters by minimizing the criterion

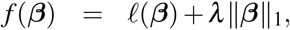

where *ℓ*(***β***) is a convex loss, *λ* is a sparsity tuning constant, and ||*β*||_1_ = Σ_*j*_|*β_j_*| is the *ℓ*_1_ norm of the parameters. The lasso has the virtues of preserving convexity and driving most parameter estimates to 0. Minimization can be conducted efficiently via cyclic coordinate descent (15; 44). The magnitude of the nonzero tuning constant *λ* determines the number of predictors selected.

Despite its widespread use, the lasso penalty has some drawbacks. First, the *ℓ*_1_ penalty tends to shrink parameters toward 0, sometimes severely so. Second, *λ* must be tuned to achieve a given model size. Third, *λ* is chosen by cross-validation, a costly procedure. Fourth and most importantly, the shrinkage caused by the penalty leaves a lot of unexplained trait variance, which tends to encourage too many false positives to enter the model ultimately identified by cross-validation.

Inflated false positive rates can be mitigated by substituting nonconvex penalties for the *ℓ*_1_ penalty. For example, the minimax concave penalty (MCP) (50)

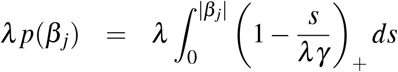

starts out at *β_j_* = 0 with slope *λ* and gradually transitions to a slope of 0 at *β_j_* = *λγ*. With minor adjustments, the coordinate descent algorithm for the lasso carries over to MCP penalized regression (8; 29). Model selection is achieved without severe shrinkage, and inference in GWAS improves (21). However, in our experience its false negative rate is considerably higher than IHT’s rate (22). A second remedy for the lasso, stability selection, weeds out false positives by looking for consistent predictor selection across random halves of the data (32). However, it is known to be under-powered for GWAS compared to standard univariate selection (2).

In contrast, iterative hard thresholding (IHT) minimizes a loss *ℓ*(***β***) subject to the nonconvex sparsity constraint ||***β***||_0_ ≤ *k*, where ||***β***||_0_ counts the number of non-zero components of *β*(3; 4; 7). Figure 1 explains graphically how the *ℓ*_0_ penalty reduces the bias of the selected parameters. In the figure *λ*, *γ*, and *k* are chosen so that the same range of *β* values are sent to zero. To its detriment, the lasso penalty shrinks all *β*’s, no matter how large their absolute values. The nonconvex MCP penalty avoids shrinkage for large *β*’s but exerts shrinkage for intermediate *β*’s. IHT, which is both nonconvex and discontinuous, avoids shrinkage altogether. For GWAS, the sparsity model-size constant *k* also has a simpler and more intuitive interpretation than the lasso tuning constant *λ*. Finally, both false positive and false negative rates are well controlled. Balanced against these advantages is the loss of convexity in optimization and concomitant loss of computational efficiency. In practice, the computational barriers are surmountable and are compensated by the excellent results delivered by IHT in high-dimensional regression problems such as multiple GWAS regression.

**Figure 1:**
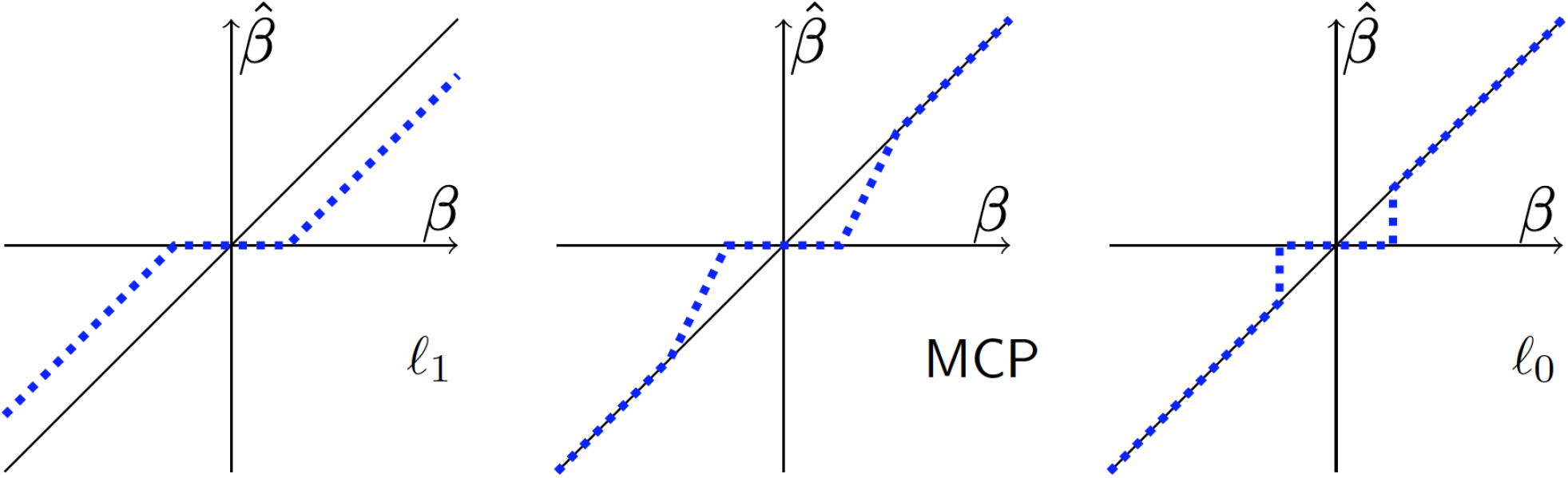
The *ℓ*_0_ quasinorm of IHT enforces sparsity without shrinkage. The estimated effect size (dashed line) is plotted against its true value (diagonal line) for *ℓ*_1_, MPC, and *ℓ*_0_ penalties.

This article has four interrelated goals. First, we extend IHT to generalized linear models. These models encompass most of applied statistics. Previous IHT algorithms focused on normal or logistic sparse regression scenarios. Our software can also perform sparse regression under Poisson and negative binomial response distributions and can be easily extended to other GLM distributions as needed. The key to our extension is the derivation of a nearly optimal step size *s* for improving the loglikelihood at each iteration. Second, we introduce doubly-sparse regression to IHT. Previous authors have considered group sparsity (46). The latter tactic limits the number of groups selected. It is also useful to limit the number of predictors selected per group. Double sparsity strikes a compromise that encourages selection of correlated causative variants in linkage disequilibrium (LD). Notably, this technique generalizes group-IHT. Third, we demonstrate how to incorporate predetermined SNP weights in IHT. Our simple and interpretable weighting option allows users to introduce prior knowledge into sparse projection. Thus, one can favor predictors whose association to the response is supported by external evidence. Fourth, we present `MendelIHT.jl:` a scalable, open source, and user friendly software for IHT in the high performance programming language Julia (6).

## 3 Model Development

This section sketches our extensions of iterative hard thresholding (IHT).

### 3.1 IHT Background

IHT was originally formulated for sparse signal reconstruction, which is framed as sparse linear least squares regression. In classical linear regression, we are given an *n* × *p* design matrix **X** and a corresponding *n*-component response vector **y**. We then postulate that **y** has mean E(**y**) = **X*β*** and that the residual vector **y** – **X*β*** has independent Gaussian components with a common variance. The parameter (regression coefficient) vector ***β*** is estimated by minimizing the sum of squares 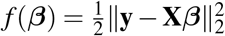. The solution to this problem is known as the ordinary least squares estimator and can be written explicitly as 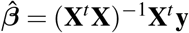, provided the problem is overdetermined (n > p). This paradigm breaks down in the high-dimensional regime *n* ≪ *p*, where the parameter vector ***β*** is underdetermined. In the spirit of parsimony, IHT seeks a sparse version of ***β*** that gives a good fit to the data. This is accomplished by minimizing *f*(***β***) subject to ||***β***||_0_ ≤ *k* for a small value of *k*, where ||·||_0_ counts the number of nonzero entries of a vector. The optimization problem is formally:

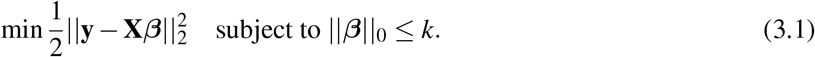

IHT abandons the explicit formula for 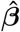 because it fails to respect sparsity and involves the numerically intractable matrix inverse (**X**^*t*^**X**)^−1^.

IHT combines three core ideas. The first is steepest descent. Elementary calculus tells us that the negative gradient −∇*f*(**x**) is the direction of steepest descent of *f*(***β***) at **x**. First-order optimization methods like IHT define the next iterate in minimization by the formula ***β***_*n*+1_ = ***β***_*n*_ + *s_n_***v**_*n*_, where **v**_*n*_ = −∇*f*(***β***_*n*_) and *s_n_* > 0 is some optimally chosen step size. In the case of linear regression −∇*f*(***β***) = **X**^*t*^(**y** – **X*β***). To reduce the error at each iteration, the optimal step size *s_n_* can be selected by minimizing the second-order Taylor expansion

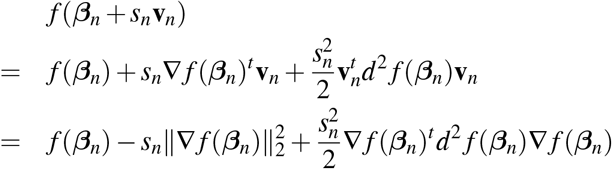

with respect to *s_n_*. Here *d*^2^ *f*(***β***) = **X**^*t*^**X** is the Hessian matrix of second partial derivatives. Because *f*(***β***) is quadratic, the expansion is exact. Its minimum occurs at the step size

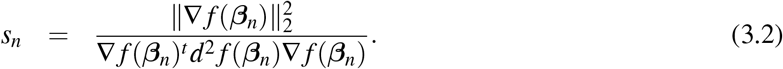

This formula summarizes the second core idea.

The third component of IHT involves projecting the steepest descent update ***β***_*n*_ + *s_n_* **v**_*n*_ onto the sparsity set *S_k_* = {***β***: ||***β***||_0_ ≤ *k*}. The relevant projection operator *P_S_k__*(***β***) sets all but the k largest entries of ***β*** in magnitude to 0. In summary, IHT solves problem (3.1) by updating the parameter vector ***β*** according to the recipe:

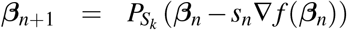

with the step size given by formula (3.2).

An optional debiasing step can be added to improve parameter estimates. This involves replacing ***β***_*n*+1_ by the exact minimum point of *f*(***β***) in the subspace defined by the support {*j*: ***β***_*n*+1, *j*_ ≠ 0} of ***β***_*n*+1_. Debiasing is efficient because it solves a low-dimensional problem. Several versions of hard-thresholding algorithms have been proposed in the signal processing literature. The first of these, NIHT (7), omits debaising. The rest, HTP(14), GraHTP (47), and CoSaMp (34) offer debiasing.

### 3.2 IHT for Generalized Linear Models

A generalized linear model (GLM) involves responses *y* following a natural exponential distribution with density in the canonical form

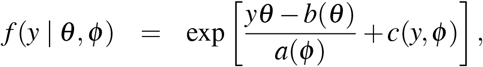

where *y* is the data, *θ* is the natural parameter, *ϕ* > 0 is the scale (dispersion), and *a*(*ϕ*), *b*(*θ*), and *c*(*y*, *ϕ*) are known functions which vary depending on the distribution (13; 30). Simple calculations show that *y* has mean *μ* = *b*′(*θ*) and variance *σ*^2^ = *b*″(*θ*)*a*(*ϕ*); accordingly, *σ*^2^ is a function of *μ*. Table 1 summarizes the mean domains and variances of a few common exponential families. Covariates enter GLM modeling through an inverse link representation *μ* = *g*(**x**^*t*^ ***β***), where **x** is a vector of covariates (predictors) and ***β*** is vector of regression coefficients (parameters). In statistical practice, data arrive as a sample of independent responses *y*_1_,…,*y_m_* with different covariate vectors **x**_1_,…, **x**_*m*_. To put each predictor on an equal footing, each should be standardized to have mean 0 and variance 1. Including an additional intercept term is standard practice.

**Table 1:** Summary of mean domains and variances for common exponential distributions. In GLM, ***μ*** = *g*(**x**^*t*^ ***β***) denotes the mean, *s* = **x**^*t*^ ***β*** the linear responses, *g* is the inverse link function, and *ϕ* the dispersion. Except for the negative binomial, all inverse links are canonical.

| Family | Mean Domain | Var(y) | $g(s)$ |
| --- | --- | --- | --- |
| Normal | $\mathbb{R}$ | $\phi^2$ | 1 |
| Poisson | $[0, \infty)$ | $\mu$ | $e^s$ |
| Bernoulli | $[0, 1]$ | $\mu(1 - \mu)$ | $\frac{e^s}{1+e^s}$ |
| Gamma | $[0, \infty)$ | $\mu^2 \phi$ | $s^{-1}$ |
| Inverse Gaussian | $[0, \infty)$ | $\mu^3 \phi$ | $s^{-1/2}$ |
| Negative Binomial | $[0, \infty)$ | $\mu(\mu \phi + 1)$ | $e^s$ |

If we assemble a design matrix **X** by stacking the row vectors 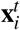, then we can calculate the loglikelihood, score, and expected information (13; 23; 30; 45)

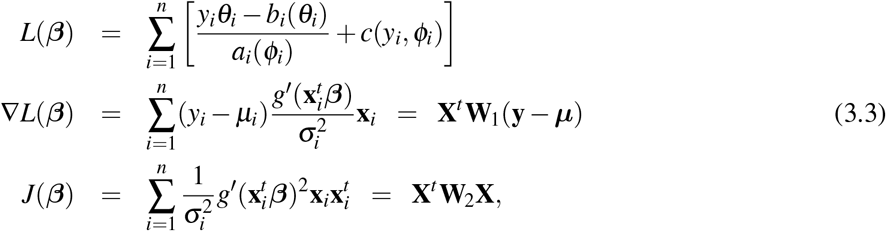

where **W**_1_ and **W**_2_ are two diagonal matrices. The second has positive diagonal entries; they coincide under the identity inverse link *g*(*s*) = *s*.

In the generalized linear model version of IHT, we maximize *L*(*β*) (equivalent to minimizing *f*(***β***) = −*L*(***β***)) and substitute the expected information *J*(***β***_*n*_) = E[−*d*^2^*L*(***β***_*n*_)] for *d*^2^*f*(***β***_*n*_) in formula (3.2). This translates into the following step size in GLM estimation:

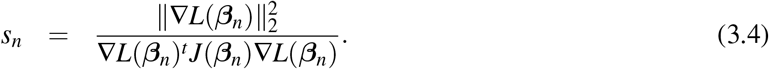

This substitution is a key ingredient of our extended IHT. It simplifies computations and guarantees that the step size is nonnegative.

### 3.3 Doubly Sparse Projections

The effectiveness of group sparsity in penalized regression has been demonstrated in general (31; 16) and for GWAS (53) in particular. Group IHT (46) enforces group sparsity but does not enforce within-group sparsity. In GWAS, model selection is desired within groups as well to pinpoint causal SNPs. Furthermore, one concern in GWAS is that two causative SNPs can be highly correlated with each other due to linkage disequilibrium (LD). When sensible group information is available, doubly sparse IHT encourages the detection of causative yet correlated SNPs while enforcing sparsity within groups. Here we discuss how to carry out a doubly-sparse projection that enforces both within- and between-group sparsity.

Suppose we divide the SNPs of a study into a collection *G* of nonoverlapping groups. Given a parameter vector ***β*** and a group *g* ∈ *G*, let ***β***_*g*_ denote the components of ***β*** corresponding to the SNPs in *g*. Now suppose we want to select at most *j* groups and at most 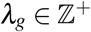 SNPs for each group *g*. In projecting ***β***, the component *β_i_* is untouched for a selected SNP *i*. For an unselected SNP, *β_i_* is reset to 0. By analogy with our earlier discussion, we can define a sparsity projection operator *P_g_*(***β***_*g*_) for each group *g*; *P_g_*(***β***_*g*_) selects the *λ_g_* most prominent SNPs in group *g*. The potential reduction in the squared distance offered by group *g* is 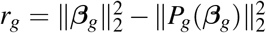. The *j* selected groups are determined by selecting the *j* largest values of *r_g_*. If desired, we can set the sparsity level *λ_g_* for each group high enough so that all SNPs in group *g* come into play. Thus, doubly-sparse IHT generalizes group-IHT. In Algorithm 1, we write *P*(***β***) for the overall projection with the component projections *P_g_*(***β***_*g*_) on the *j* selected groups and projection to zero on the remaining groups.

### 3.4 Prior weights in IHT

Zhou et al. (53) treat prior weights in penalized GWAS. Before calculating the lasso penalty, they multiply each component of the parameter vector ***β*** by a positive weight *w_i_*. We can do the same in IHT before projection. Thus, instead of projecting the steepest descent step ***β*** = ***β***_*n*_ + *s_n_* **v**_*n*_, we project the Hadamard (pointwise) product **w** ○ ***β*** of ***β*** with a weight vector **w**. This produces a vector with a sensible support *S*. The next iterate ***β***_*n*+1_ is defined to have support *S* and to be equal to ***β***_*n*_ + *s_n_* **v**_*n*_ on *S*.

In GWAS, weights can and should be informed by prior biological knowledge. A simple scheme for choosing nonconstant weights relies on minor allele frequencies. For instance, Zhou et al. (51) assign SNP *i* with minor allele frequency *p_i_* the weight 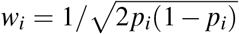. Giving rare SNPs greater weight in this fashion is most appropriate for traits under strong negative selection (49; 37). Alternatively, our software permits users to assign weights geared to specific pathway and gene information.

de Lamare et al. (12) incorporate prior weights into IHT by adding an element-wise logarithm of a weight vector **q** before projection. The weight vector **q** is updated iteratively and requires two additional tuning constants that in practice are only obtained through cross validation. Our weighting scheme is simpler, more computationally efficient, and more interpretable.

### 3.5 Algorithm Summary

The final algorithm combining doubly sparse projections, prior weight scaling, and debiasing is summarized in Algorithm 1.

**Algorithm 1.**
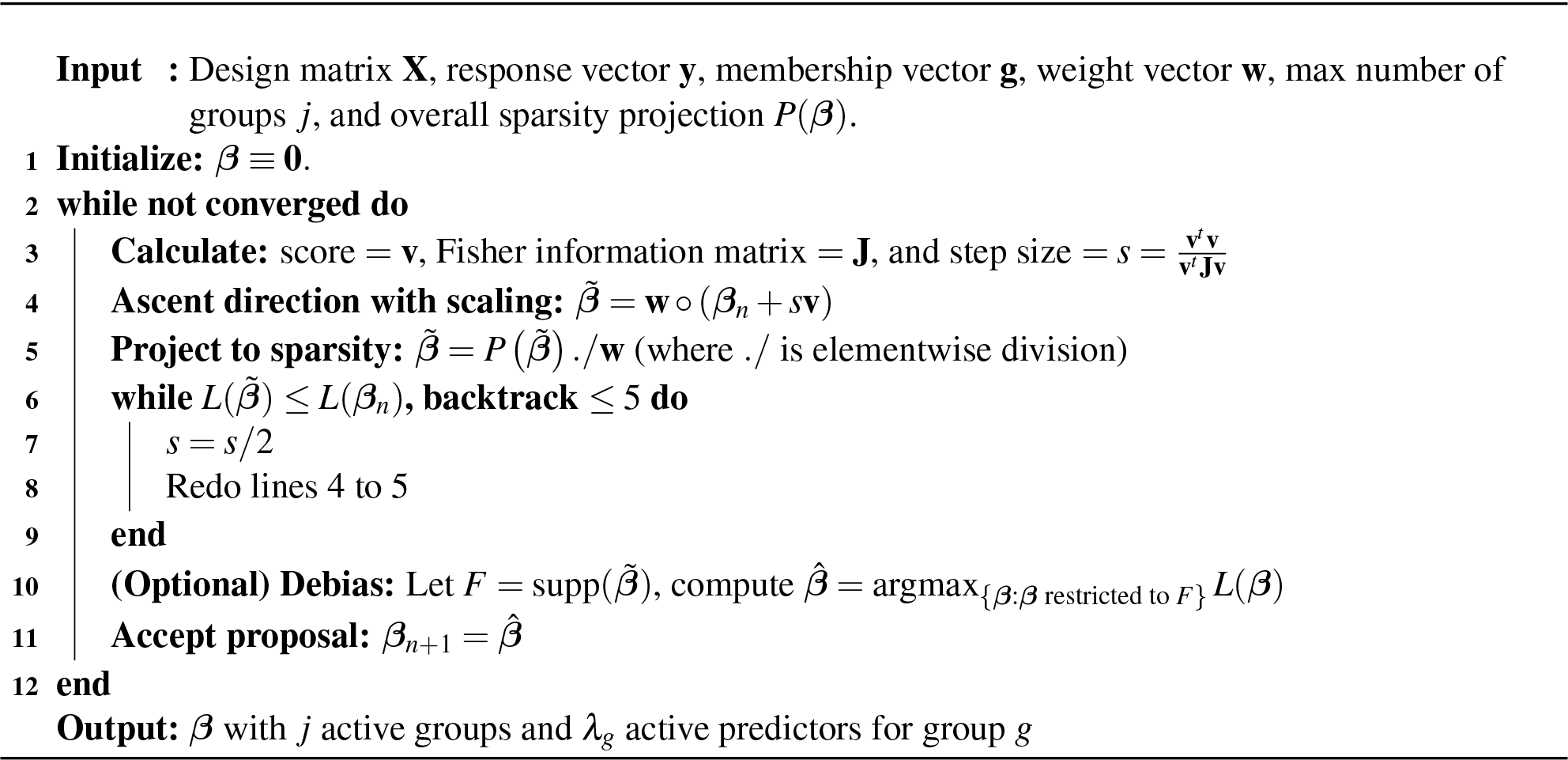
Iterative hard-thresholding.

## 4 Results

Readers can reproduce our results by accessing the software, documentation, and Jupyter notebooks at: https://github.com/OpenMendel/MendelIHT.jl

### 4.1 Scalability of IHT

To test the scalability of our implementation, we ran IHT on *p* = 10^6^ SNPs for sample sizes *n* = 10,000,20,000,…, 120,000 with five independent replicates per *n*. All simulations rely on a true sparsity level of *k* = 10. Based on an Intel-E5-2670 machine with 63GB of RAM and a single 3.3GHz processor, Figure 2 plots the IHT median CPU time per iteration, median iterations to convergence, and median memory usage under Gaussian, logistic, Poisson, and negative binomial models. The largest matrix simulated here is 30GB in size and can still fit into our personal computer’s memory. Of course, it is possible to test even larger sample sizes using cloud or cluster resources, which are often needed in practice.

**Figure 2:**
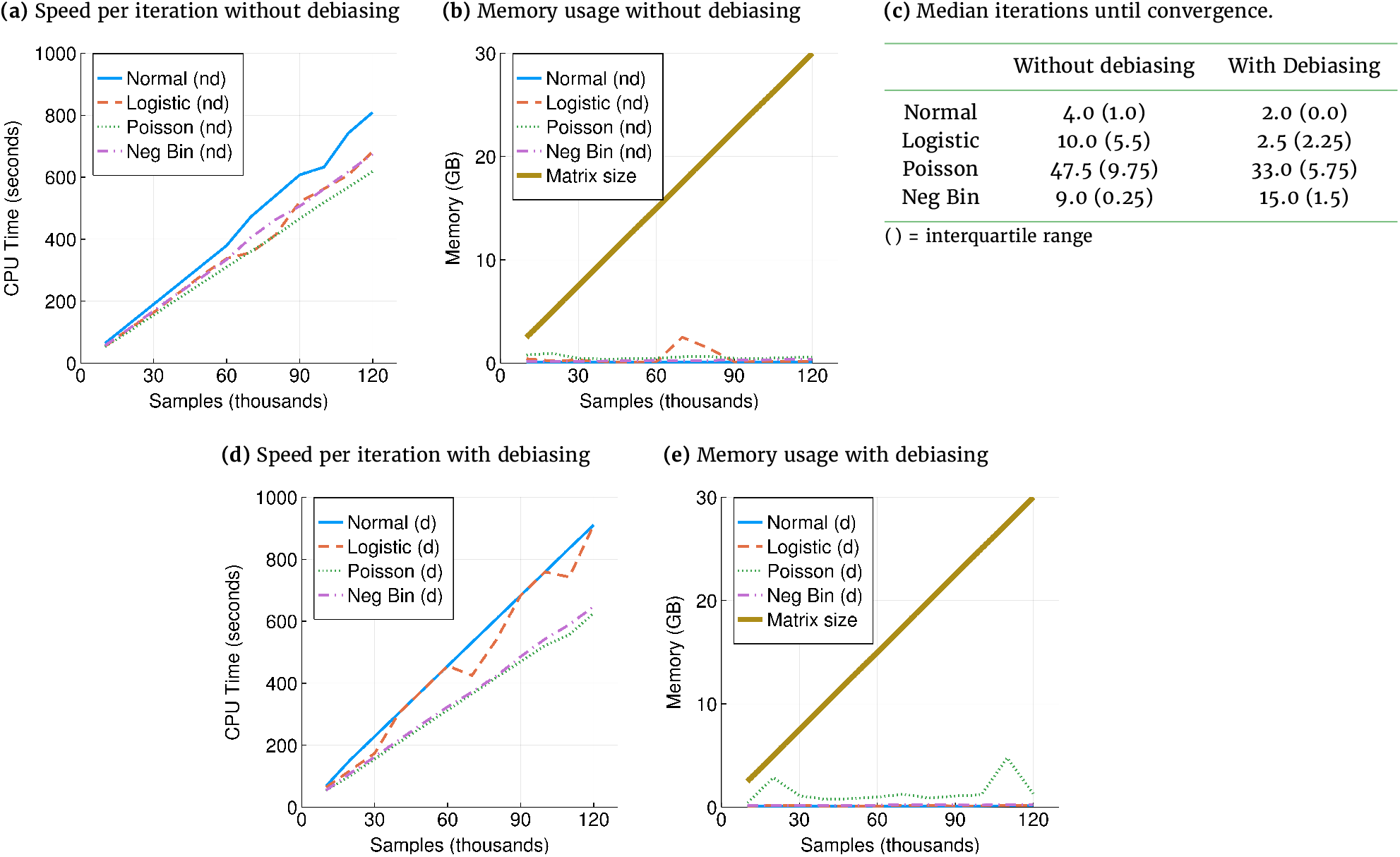
(a, d) Time per iteration scales linearly with data size. Speed is measured for compressed genotype files. On uncompressed data, all responses are roughly 10 times faster. (b, e) Memory usage scales as ~ 2*np* bits. Note memory for each response are usages in addition to loading the genotype matrix. Uncompressed data requires 32 times more memory. (c) Debiasing reduces median iterations until convergence for all but negative binomial regression. Benchmarks were carried out on 10^6^ SNPs and sample sizes ranging from 10,000 to 120,000. Hence, the largest matrix here requires 30GB and can still fit into personal computer memories.

The formation of the vector ***μ*** of predicted values requires only a limited number of nonzero regression coefficients. Consequently, the computational complexity of this phase of IHT is relatively light. In contrast, calculation of the Fisher score (gradient) and information (expected negative Hessian) depend on the entire genotype matrix **X**. Fortunately, each of the *np* entries of **X** can be compressed to 2 bits. Figure 2b and d show that IHT memory demands beyond storing **X** never exceeded a few gigabytes. Figure 2a and c show that IHT run time per iteration increases linearly in problem size *n*. Similarly, we expect increasing *p* will increase run time linearly, since the bottleneck of IHT is the matrix-vector multiplication step in computing the gradient, which scales as *O*(*np*). Debiasing increases run time per iteration only slightly. Except for negative binomial responses, debiasing is effective in reducing the number of iterations required for convergence and hence overall run time.

### 4.2 Cross Validation in Model Selection

In actual studies, the true number of genetic predictors *k*_true_ is unknown. This section investigates how *q*-fold cross-validation can determine the best model size on simulated data. Under normal, logistic, Poisson, and negative binomial models, we considered 50 different combinations of **X, y**, and ***β***_true_ with *k*_true_ = 10, *n* = 5000 samples, and *p* = 50,000 SNPs fixed in all replicates. Here, *k*_true_ is chosen so that it is closer to our NFBC and UK Biobank results. On these data sets we conducted 5-fold cross validation across 20 model sizes *k* ranging from 1 to 20. Figure 3 plots deviance residuals on the holdout dataset for each of the four GLM responses (mean squared error in the case of normal responses) and the best estimate 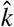 of *k*_true_.

**Figure 3:**
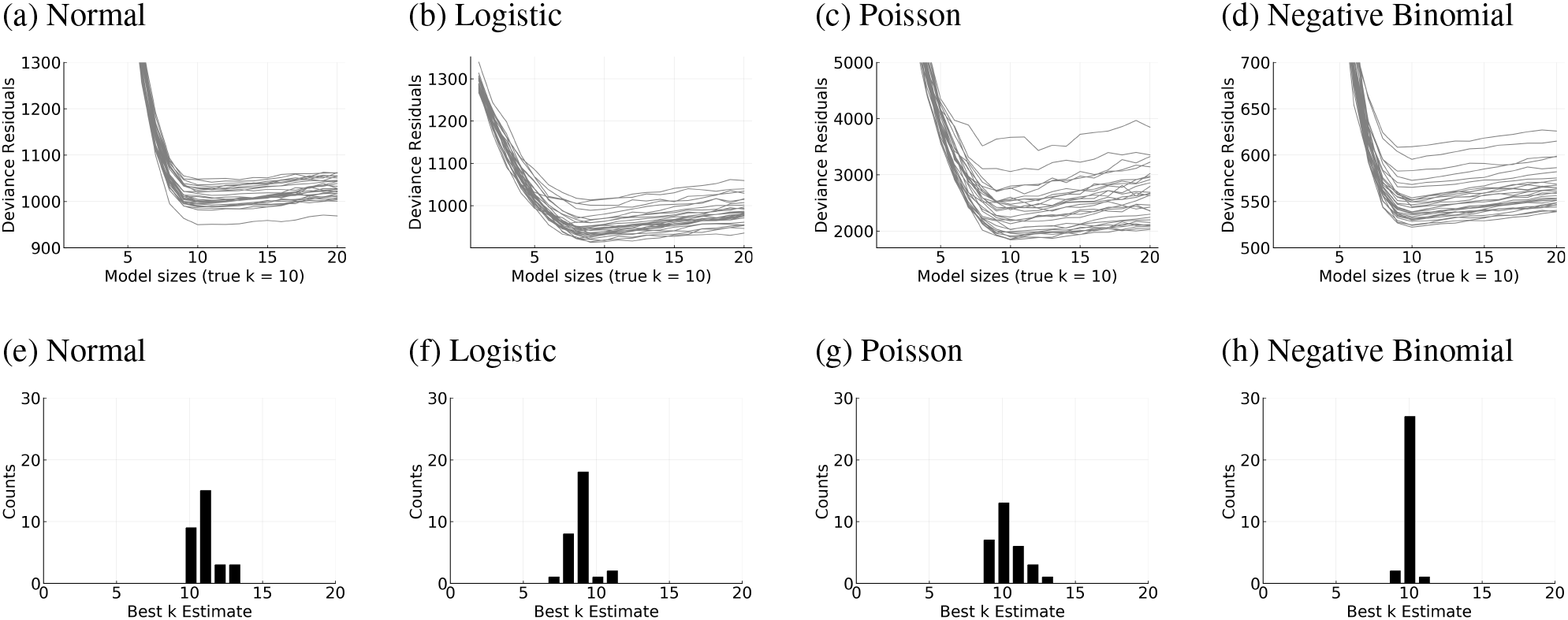
Five-fold cross validation results is capable of identifying the true model size *k*_true_. (a-d) Deviance residuals of the testing set are minimized when the estimated model size 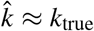. Each line represents 1 simulation. (e-h) 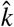 is narrowly spread around *k*_true_ = 10.

Figure 3 shows that *k*_true_ can be effectively recovered by cross validation. In general, prediction error starts off high where the proposed sparsity level *k* severely underestimates *k*_true_ and plateaus when *k*_true_ is reached (Figure 3a-d). Furthermore, the estimated sparsity 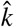 for each run is narrowly centered around *k*_true_ = 10 (Figure 3e-f). In fact, 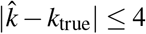 always holds. When 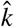 exceeds *k*_true_, the estimated regression coefficients for the false predictors tend to be very small. In other words, IHT is robust to overfitting, in contrast to lasso penalized regression. We see qualitatively similar results when *k*_true_ is large. This proved to be the case in our previous paper (22) for Gaussian models with *k*_true_ ∈ {100,200,300}.

### 4.3 Comparing IHT to Lasso and Marginal Tests in Model Selection

Comparison of the true positive and false positive rates of IHT and its main competitors is revealing. For lasso regression we use the `glmnet` implementation of cyclic coordinate descent (15; 43; 44) (v2.0-16 implemented in `R 3.5.2`); for marginal testing we use the beta version of `MendelGWAS` (54). As explained later, Poisson regression is supplemented by zero-inflated Poisson regression implemented under the `pscl` (48) (v1.5.2) package of `R`. Unfortunately, `glmnet` does not accommodate negative binomial regression. Because both `glmnet` and `pscl` operate on floating point numbers, we limit our comparisons to small problems with 1000 subjects, 10,000 SNPs, 50 replicates, and *k* = 10 causal SNPs. IHT performs model selection by 3-fold cross validation across model sizes ranging from 1 to 50. This range is generous enough to cover the models selected by lasso regression. We adjust for multiple testing in the marginal case test by applying a p-value cutoff of 5 × 10^−6^.

Table 2 demonstrates that IHT achieves the best balance between maximizing true positives and minimizing false positives. IHT finds more true positives than marginal testing and almost as many as lasso regression. IHT also finds far fewer false positives than lasso regression. Poisson regression is exceptional in yielding an excessive number of false positives in marginal testing. A similar but less extreme trend is observed for lasso regression. The marginal false positive rate is reduced by switching to zero-inflated Poisson regression. This alternative model is capable of handling overdispersion due an excess of 0 values. Interestingly, IHT rescues the Poisson model by accurately capturing the simultaneous impact of multiple predictors.

**Table 2:** IHT achieves the best balance of false positives and true positives compared to lasso and marginal (single-snp) regression. TP = true positives, FP = false positives. There are *k* = 10 causal SNPs. Best model size for IHT and lasso were chosen by cross validation. () = zero-inflated Poisson regression.

|  | Normal | Logistic | Poisson | Neg Bin |
| --- | --- | --- | --- | --- |
| IHT TP | 8.84 | 6.28 | 7.2 | 9.0 |
| IHT FP | 0.02 | 0.1 | 1.28 | 0.98 |
| Lasso TP | 9.52 | 8.16 | 9.28 | NA |
| Lasso FP | 31.26 | 45.76 | 102.24 | NA |
| Marginal TP | 7.18 | 5.76 | 9.04 (5.94) | 5.98 |
| Marginal FP | 0.06 | 0.02 | 1527.9 (0.0) | 0.0 |

### 4.4 Reconstruction Quality for GWAS Data

Table 3 demonstrates that IHT estimates show little bias compared to estimates from lasso and marginal regressions. These trends hold with or without debiasing as described earlier. The proportion of variance explained is approximately the same in both scenarios. The displayed values are the averaged estimated *β*’s, computed among the SNPs actually found. As expected, lasso estimates show severe shrinkage compared to IHT. Estimates from marginal tests are severely overestimated, since each SNP are asked to explain more trait variance than it could. As the magnitude of ***β**_true_* falls, IHT estimates show an upward absolute bias, consistent with the winner’s curse phenomenon. When sample sizes are small, small effect sizes make most predictors imperceptible amid the random noise. The winner’s curse operates in this regime and cannot be eliminated by IHT. Lasso’s strong shrinkage overwhelms the bias of the winner’s curse and yields estimates smaller than true values.

The results displayed in Table 3 reflect *n* = 5, 000 subjects, *p* = 10,000 SNPs, 100 replicates, and a sparsity level *k* fixed at its true value *k*_true_ = 10. The *λ* value for lasso is chosen by cross validation. To avoid data sets with monomorphic SNPs, the minimum minor allele frequency (maf) is set at 0.05. For linear, logistic and Poisson regressions in marginal tests, we first screen for potential SNPs via a score test. Only top SNPs are used in the more rigorous and more computationally intensive likelihood ratio tests, which gives the beta estimates. This procedure is described in (54). We ran likelihood ratio tests for all SNPs in the negative binomial model because the screening procedure is not yet implemented. However, the inflation in parameter estiamtes are present throughout all marginal tests.

**Table 3:** Comparison of coefficient estimates among IHT, lasso, and marginal regression methods. Displayed coefficients are average fitted valued ± one standard error for the discovered predictors. * = zero true positives observed on average. NA = `glmnet` does not support negative binomial lasso regression. There are *k* = 10 true SNPs.

| $\beta_{\text{true}}$ | $\beta_{\text{IHT}}^{\text{Normal}}$ | $\beta_{\text{IHT}}^{\text{Logistic}}$ | $\beta_{\text{IHT}}^{\text{Poisson}}$ | $\beta_{\text{IHT}}^{\text{NegBin}}$ |
| --- | --- | --- | --- | --- |
| .5 | .501 ± .015 | .508 ± .039 | .492 ± .039 | .567 ± .670 |
| .25 | .249 ± .013 | .256 ± .038 | .247 ± .012 | .249 ± .012 |
| .10 | .097 ± .014 | .125 ± .016 | .100 ± .014 | .010 ± .012 |
| .05 | .063 ± .007 | .108 ± .006 | .057 ± .008 | .060 ± .008 |
| $\beta_{\text{true}}$ | $\beta_{\text{lasso}}^{\text{Normal}}$ | $\beta_{\text{lasso}}^{\text{Logistic}}$ | $\beta_{\text{lasso}}^{\text{Poisson}}$ | $\beta_{\text{lasso}}^{\text{NegBin}}$ |
| .5 | .451 ± .015 | .366 ± .058 | .458 ± .037 | NA |
| .25 | .199 ± .013 | .137 ± .032 | .208 ± .015 | NA |
| .10 | .046 ± .014 | .022 ± .016 | .058 ± .016 | NA |
| .05 | .012 ± .008 | .008 ± .003 | .012 ± .009 | NA |
| $\beta_{\text{true}}$ | $\beta_{\text{marginal}}^{\text{Normal}}$ | $\beta_{\text{marginal}}^{\text{Logistic}}$ | $\beta_{\text{marginal}}^{\text{Poisson}}$ | $\beta_{\text{marginal}}^{\text{NegBin}}$ |
| .5 | .990 ± .500 | .983 ± .475 | .942 ± .331 | .930 ± .315 |
| .25 | .493 ± .189 | .480 ± .216 | .452 ± .184 | .486 ± .178 |
| .10 | .203 ± .078 | * | .198 ± .097 | .190 ± .090 |
| .05 | * | * | .165 ± .049 | .097 ± .060 |

### 4.5 Correlated Covariates and Doubly Sparse Projections

Next we study how well IHT works on correlated data and whether doubly-sparse projection can enhance model selection. Table 4 shows that, in the presence of extensive LD, IHT performs reasonably well even without grouping information. When grouping information is available, group IHT enhances model selection. The results displayed in Table 4 reflect *n* = 1,000 samples, *p* = 10,000 SNPs, and 100 replicates. Each SNP belongs to 1 of 500 disjoint groups containing 20 SNPs each; *j* = 5 distinct groups are each assigned 1,2,…,5 causal SNPs with effect sizes randomly chosen from {−0.2,0.2}. In all there 15 causal SNPs. For grouped-IHT, we assume perfect group information. That is, groups containing 1 ~ 5 causative SNPs are assigned *λ_g_* ∈ {1,2,…,5}. The remaining groups are assigned *λ_g_* = 1. As described in the Methods Section, the simulated data show LD within each group, with the degree of LD between two SNPs decreasing as their separation increases. Although the conditions of this simulation are somewhat idealized, they mimic what might be observed if small genetic regions of whole exome data were used with IHT.

**Table 4:** Doubly-sparse IHT enhances model selection on simulated data. TP = true positives, FP = false positives, ± 1 standard error. There are 15 causal SNPs in 5 groups, each containing *k* ∈ ={1,2,…5} SNPs.

|  | Ungrouped-IHT |  | Grouped-IHT |  |
| --- | --- | --- | --- | --- |
|  | TP | FP | TP | FP |
| Normal | 11.1 ± 1.9 | 3.9 ± 1.9 | 12.2 ± 2.0 | 2.8 ± 2.0 |
| Logistic | 3.8 ± 1.6 | 11.2 ± 1.6 | 7.7 ± 2.2 | 7.3 ± 2.2 |
| Poisson | 11.5 ± 2.2 | 3.5 ± 2.2 | 12.4 ± 1.7 | 2.6 ± 1.7 |
| Neg Bin | 11.0 ± 2.1 | 4.0 ± 2.1 | 12.4 ± 1.6 | 2.6 ± 1.6 |

We repeated this examination of doubly sparse projection for the first 30,000 SNPs of the NFBC1966 (36) data for all samples passing the quality control measures outlined in our Methods Section. We arbitrarily assembled 2 large groups with 2000 SNPs, 5 medium groups with 500 SNPs, and 10 small groups with 100 SNPs, representing genes of different length. The remaining SNPs are lumped into a final group representing non-coding regions. In all there are 18 groups. Since group assignments are essentially random beyond choosing neighboring SNPs, this example represents the worse case scenario of a relatively sparse marker map with undifferentiated SNP groups. We randomly selected 1 large group, 2 medium groups, and 3 small groups to contain 5, 3, and 2 causal SNPs, respectively. The non-coding region harbors 2 causal SNPs. In all there are 19 causal SNPs. Effect sizes were randomly chosen to be −0.2 or 0.2. We ran 100 independent simulation studies under this setup, where the large, medium, small, and non-coding groups are each allowed 5, 3, 2, and 2 active SNPs. The results are displayed in Table 5. We find that even in this worse case scenario where group information is completely lacking that grouped IHT does no worse than ungrouped IHT.

**Table 5:**
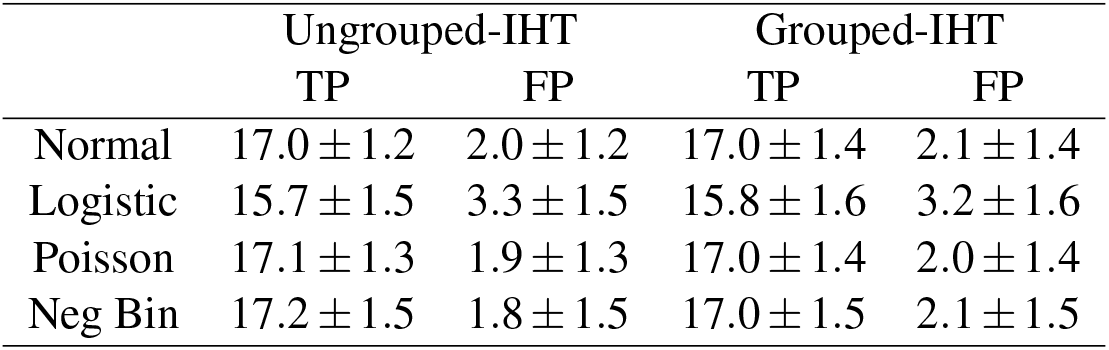
Doubly sparse IHT is comparable to regular IHT on NFBC dataset using arbitrary groups. TP = true positives, FP = false positives, ± 1 standard error. There are 19 causal SNPs in 18 groups of various size. Simulation was carried out on the first 30,000 SNPs of the NFBC1966 (36) dataset.

### 4.6 Introduction of Prior Weights

This section considers how scaling by prior weights helps in model selection. Table 6 compares weighted IHT reconstructions with unweighted reconstructions where all weights *w_i_* = 1. The weighted version of IHT consistently finds approximately 10% more true predictors than the unweighted version. Here we simulated 50 replicates involving 1000 subjects, 10,000 uncorrelated variants, and *k* = 10 true predictors for each GLM. For the sake of simplicity, we defined a prior weight *w_i_* = 2 for about one-tenth of all variants, including the 10 true predictors. For the remaining SNPs the prior weight is *w_i_* = 1. These choices reflect a scenario where one tenth of all genotyped variants fall in a protein coding region, including the 10 true predictors, and where such variants are twice as likely to influence a trait as those falling in non-coding regions.

**Table 6:**
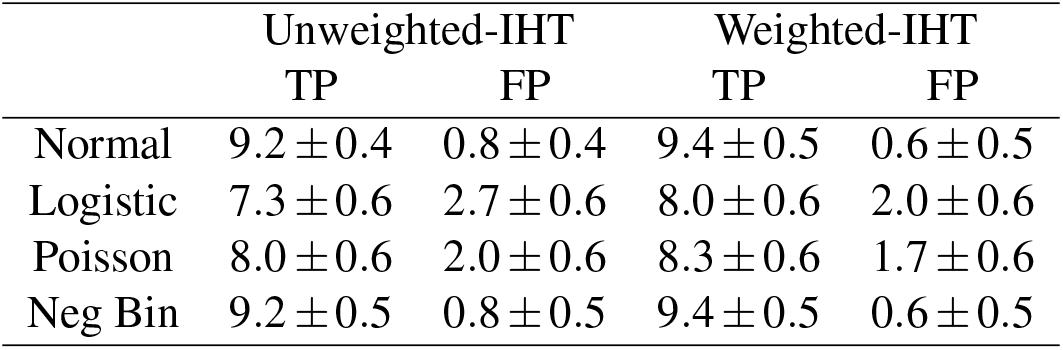
Weighted IHT enhances model selection. TP = true positives, FP = false positives, ± 1 standard error. The true number of SNPs is *k* = 10.

|  | Unweighted-IHT |  | Weighted-IHT |  |
| --- | --- | --- | --- | --- |
|  | TP | FP | TP | FP |
| Normal | $9.2 \pm 0.4$ | $0.8 \pm 0.4$ | $9.4 \pm 0.5$ | $0.6 \pm 0.5$ |
| Logistic | $7.3 \pm 0.6$ | $2.7 \pm 0.6$ | $8.0 \pm 0.6$ | $2.0 \pm 0.6$ |
| Poisson | $8.0 \pm 0.6$ | $2.0 \pm 0.6$ | $8.3 \pm 0.6$ | $1.7 \pm 0.6$ |
| Neg Bin | $9.2 \pm 0.5$ | $0.8 \pm 0.5$ | $9.4 \pm 0.5$ | $0.6 \pm 0.5$ |

### 4.7 Hypertension GWAS in the UK Biobank

Now we test IHT on the second release of UK Biobank (38) data. This dataset contains ~ 500,000 samples and ~ 800,000 SNPs without imputation. Phenotypes are systolic blood pressure (SBP) and diastolic blood pressure (DBP), averaged over 4 or fewer readings. To adjust for ancestry and relatedness, we included the following nongenetic covariates: sex, hospital center, age, age^2^, BMI, and the top 10 principal components computed with `FlashPCA2` (1). After various quality control procedures as outlined in the Methods section, the final dataset used in our analysis contains 185,565 samples and 470,228 SNPs. For UK biobank analysis, we omitted debiasing, prior weighting, and doubly sparse projections.

#### 4.7.1 Stage 2 Hypertension under a Logistic Model

Consistent with the clinical definition for stage 2 hypertension (S2 Hyp) (42), we designated patients as hypertensive if their SBP ≥ 140mmHG or DBP ≥ 90 mmHG. We ran 5-fold cross validated logistic model across model sizes *k* = {1,2,…, 50}. The work load was distributed to 50 computers, each with 5 CPU cores. Each computer was assigned one model size, and all completed its task within 24 hours. The model size that minimizes the deviance residuals is 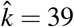. The selected predictors include the 33 SNPs listed in Table 7 and 6 non-genetic covariates: intercept, sex, age, age^2^, BMI, and the fifth principal component.

**Table 7:** UK Biobank GWAS results generated by running IHT on Stage 2 Hypertension (S2 Hyp) under a logistic model. The SNP ID, chromosome number, position (in basepair), and estimated effect sizes are listed.

| SNP | Chrom | Position | $\hat{\beta}$ | Known? |
| --- | --- | --- | --- | --- |
| rs17367504 | 1 | 11862778 | 0.046 | (27; 18) |
| rs757110 | 2 | 17418477 | -0.025 |  |
| rs1898841 | 2 | 165070207 | 0.022 | (18) |
| rs1374264 | 2 | 164999883 | 0.020 | (18) |
| rs16998073 | 4 | 81184341 | -0.048 | (27; 18) |
| rs1173771 | 4 | 32815028 | 0.046 | (27; 18) |
| rs13107325 | 4 | 103188709 | 0.030 | (27; 18) |
| rs72742749 | 5 | 32834974 | 0.029 |  |
| rs11241955 | 5 | 127626884 | 0.028 |  |
| rs2072495 | 5 | 158296996 | -0.027 |  |
| rs805293 | 6 | 31688518 | -0.029 | (18) |
| rs2392929 | 7 | 106414069 | -0.039 | (27; 18) |
| rs73203495 | 8 | 11580334 | -0.031 |  |
| rs12258967 | 10 | 18727959 | 0.039 | (27; 18) |
| rs11191580 | 10 | 104906211 | 0.039 | (27; 18) |
| rs2274224 | 10 | 96039597 | 0.036 | (18) |
| rs1530440 | 10 | 63524591 | 0.028 | (27; 18) |
| rs10895001 | 11 | 100533021 | 0.043 | (18) |
| rs2293579 | 11 | 47440758 | -0.035 | (18) |
| rs2923089 | 11 | 10357572 | -0.029 | (18) |
| rs762551 | 11 | 75041917 | -0.027 | (18) |
| rs4548577 | 11 | 46998512 | 0.026 |  |
| rs2681492 | 12 | 90013089 | 0.030 | (27; 18) |
| rs10849937 | 12 | 111792427 | 0.030 | (18) |
| rs35085068 | 14 | 23409909 | -0.027 | (18) |
| rs12901664 | 15 | 98338524 | -0.027 |  |
| rs7497304 | 15 | 91429176 | -0.021 | (27; 18) |
| rs2677738 | 15 | 91441673 | 0.021 | (18) |
| rs3744760 | 17 | 43195981 | -0.043 | (18) |
| rs292445 | 18 | 55897720 | -0.026 |  |
| rs167479 | 19 | 11526765 | 0.036 | (27; 18) |
| rs34328549 | 19 | 7253184 | 0.035 | (18) |
| rs16982520 | 20 | 57758720 | -0.030 | (27; 18) |

Figure 4, generated by `MendelPlots.jl` (19), compares univariate logistic GWAS with logistic IHT. SNPs recovered by IHT are circled in black. Our Github page records the full list of significant SNPs detected by univariate GWAS. There are 10 SNPs selected by IHT that have a p-value less than 5 × 10^−8^; 83 SNPs pass the threshold in the univariate analysis but remain unselected by IHT. IHT tends to pick the most significant SNP among a group of SNPs in LD. Table 7 shows 25 SNPs selected by IHT that were previously reported to be associated with elevated SBP/DBP (27) or that exhibit genome-wide significance when the same data are analyzed as an ordinal trait (18). Ordinal univariate GWAS treats the different stages of hypertension as ordered categories. Ordinal GWAS has higher power than logistic or multinomial GWAS (18). The known SNPs displayed in Table 7 tend to have larger absolute effect sizes (avg 0.033) than the unknown SNPs (avg = 0.027). Finally, IHT is able to recover two pairs of highly correlated SNPs: (rs1374264, rs1898841) and (rs7497304, rs2677738) with pairwise correlations of *r*_1,2_ = 0.59 and *r*_3,4_ = 0.49.

**Figure 4:**
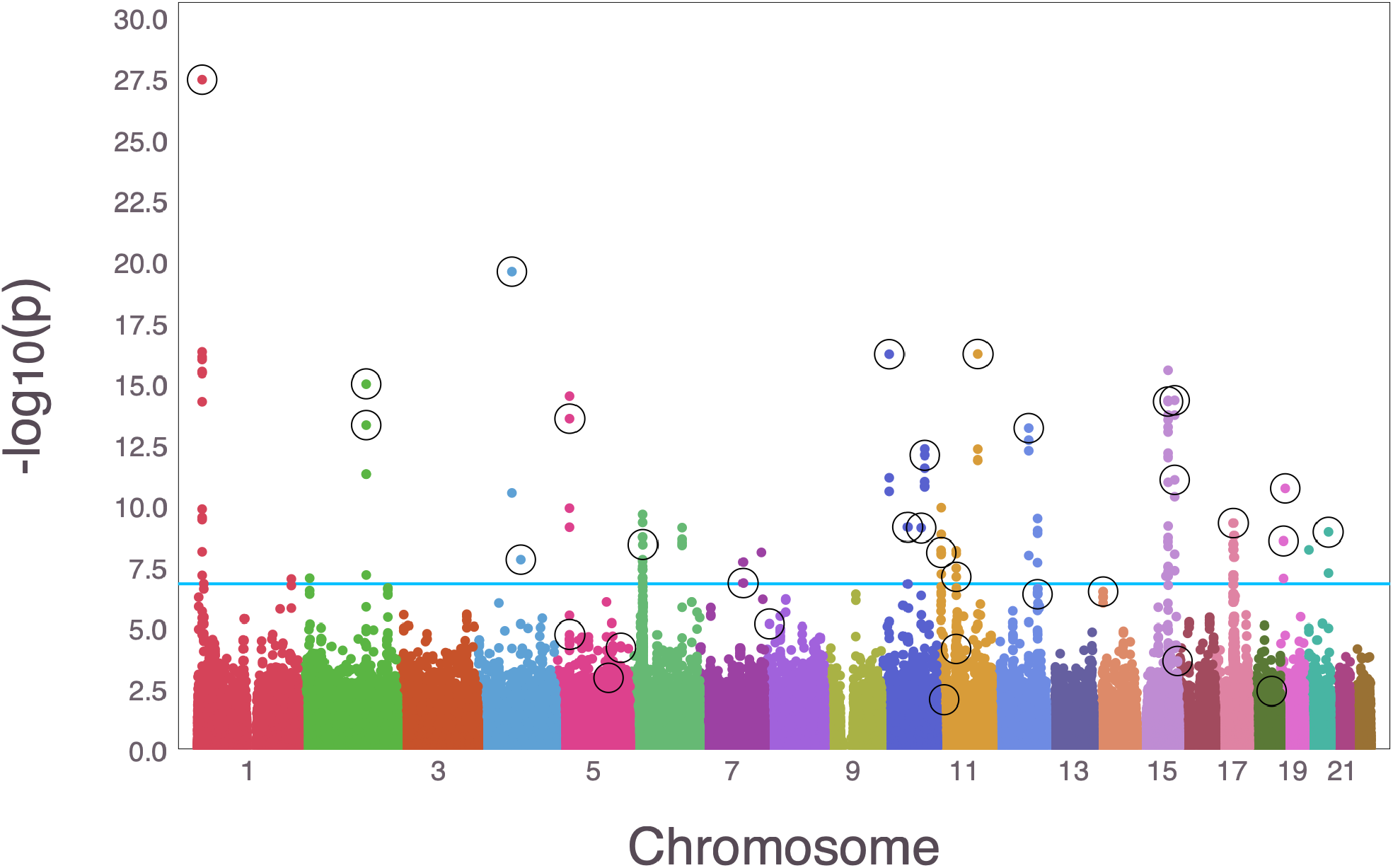
Manhattan plot comparing a logistic (univariate) GWAS vs logistic IHT on UK Biobank data. Colored dots are log_10_ p-values from a logistic GWAS, and the circled dots are SNPs recovered by IHT.

### 4.8 Cardiovascular GWAS in NFBC1966

We also tested IHT on data from the 1966 Northern Finland Birth Cohort (NFBC1966) (36). Although this dataset is relatively modest with 5402 participants and 364,590 SNPs, it has two virtues. First, it has been analyzed multiple times (22; 36; 17), so comparison with earlier analysis is easy. Second, due to a population bottleneck (28), the participants’ chromosomes exhibit more extensive linkage disequilibrium than is typically found in less isolated populations. Multiple regression methods, including the lasso, have been criticized for their inability to deal with the dependence among predictors induced by LD. Therefore this dataset provides an interesting test case.

#### 4.8.1 High Density Lipoprotein (HDL) Phenotype as a Normal model

Using IHT we find previously associated SNPs as well as a few new potential associations. We model the HDL phenotype as normally-distributed and find a best model size 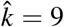 based on 5-fold cross validation across model sizes *k* = {1,2,…, 20}. Without debiasing, the analysis was completed in 2 hours and 4 minutes with 30 CPU cores on a single machine. Table 8 displays the recovered predictors. SNP rs1800961 was replaced by rs7499892 with similar effect size if we add the debiasing step in obtaining the final model.

**Table 8:** NFBC GWAS results generated by running IHT on high density lipoprotein (HDL) phenotype as a normal response. The SNP ID, chromosome number, position (in basepair), and estimated effect sizes are listed.

| SNP | Chrom | Position | $\hat{\beta}$ | Known? |
| --- | --- | --- | --- | --- |
| rs6917603 | 6 | 30125050 | 0.17 | (22; 27) |
| rs9261256 | 6 | 30129920 | -0.07 | (22) |
| rs9261224 | 6 | 30121866 | -0.03 |  |
| rs7120118 | 11 | 47242866 | -0.03 | (22; 36; 27) |
| rs1532085 | 15 | 56470658 | -0.04 | (22; 36; 27) |
| rs3764261 | 16 | 55550825 | -0.05 | (22; 36; 27) |
| rs3852700 | 16 | 65829359 | -0.03 |  |
| rs1800961 | 20 | 42475778 | 0.03 | (27) |

Importantly, IHT is able to simultaneously recover effects for SNPs (1) rs9261224, (2) rs6917603, and (3) rs6917603 with pairwise correlations of *r*_1,2_ = 0.618, *r*_1,3_ = 0.984, and *r*_2,3_ = 0.62. This result is achieved without grouping of SNPs, which can further increase association power. Compared with earlier analyses of these data, we find 3 SNPs that were not listed in our previous IHT paper (22), presumably due to slight algorithmic modifications. The authors of NFBC (36) found 5 SNPs associated with HDL under SNP-by-SNP testing. We did not find SNPs rs2167079 and rs255049. To date, rs255049 was replicated (17). SNP rs2167079 has been reported to be associated with an unrelated phenotype (33). If we repeat the analysis with HDL dichotomized into low and high HDL using a cutpoint of 60ml/DL, then we identify the 5 SNPs rs9261224, rs6917603, rs9261256, rs3764261, and rs9898058; all but one of these, SNP rs9898058, is also found under the continuous model. This SNP is not replicated in previous studies. As in the continuous model, rs6917603 has the largest effect of all the selected SNPs. Readers interested in the full result can visit our Github site.

#### 4.8.2 Low Density Lipoprotein (LDL) as a Binary Response

Unfortunately we did not have access to any qualitative phenotypes for this cohort, so for purposes of illustration, we fit a logistic regression model to a derived dichotomous phenotype, high versus low levels of Low Density Lipoprotein (LDL). The original data are continuous, so we choose 145 mg/dL, the midpoint (42) between the borderline-high and high LDL cholesterol categories, to separate the two categories. This dichotomization resulted in 932 cases (high LDL) and 3961 controls (low LDL). Under 5-fold cross validation without debiasing across model sizes *k* = {1,2,…,20}, we find 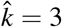. Using 30 CPU cores, our algorithm finishes in 1 hours and 7 minutes.

Despite the loss of information inherent in dichotomization, our results are comparable to the prior results under a normal model for the original quantitative LDL phenotype. Our final model still recovers two SNP predictors with and without debiasing (Table 8). We miss all but one of the SNPs that the NFBC analysis found to be associated with LDL treated as a quantitative trait. Notably we again find an association with SNP rs6917603 that they did not report.

## 5 Discussion

Multiple regression methods like iterative hard thresholding provide a principled way of model fitting and variable selection. With increasing computing power and better software, multiple regression methods are likely to prevail over univariate methods. This paper introduces a scalable implementation of iterative hard thresholding for generalized linear models. Because lasso regression can handle group and prior weights, we have also extended IHT to incorporate such prior knowledge. When it is available, enhanced IHT outperforms standard IHT. Given its sharper parameter estimates and more robust model selection, IHT is clearly superior to lasso selection or marginal association testing in GWAS.

Our real data analyses and simulation studies suggest that IHT can (a) recover highly correlated SNPs, (b) avoid over-fitting, (c) deliver better true positive and false positive rates than either marginal testing or lasso regression, (d) recover unbiased regression coefficients, and (e) exploit prior information and group-sparsity. Our Julia implementation of IHT can also exploit parallel computing strategies that scale to biobank-level data. In our opinion, the time is ripe for the genomics community to embrace multiple regression models as a supplement to and possibly a replacement of marginal analysis.

Although we focused our attention on GWAS, the potential applications of iterative hard thresholding reach far beyond gene mapping. Our IHT implementation accepts arbitrary numeric data and is suitable for a variety of applied statistics problems. Genetics and the broader field of bioinformatics are blessed with rich, ultra-high dimensional data. IHT is designed to solve such problems. By extending IHT to the realm of generalized linear models, it becomes possible to fit regression models with more exotic distributions than the Gaussian distributions implicit in ordinary linear regression. In our view IHT will eventually join and probably supplant lasso regression as the method of choice in GWAS and other high-dimensional regression settings.

## 6 Methods

### 6.1 Data Simulation

Our simulations mimic scenarios for a range of rare and common SNPs with or without LD. Unless otherwise stated, we designate 10 SNPs to be causal with effect sizes of 0.1,0.2,…, 1.0.

To generate independent SNP genotypes, we first sample a minor allele frequency *ρ_j_* ~ Uniform(0,0.5) for each SNP *j*. To construct the genotype of person *i* at SNP *j*, we then sample from a binomial distribution with success probability *ρ_j_* and two trials. The vector of genotypes (minor allele counts) for person *i* form row 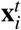 of the design matrix **X**. To generate SNP genotypes with linkage disequilibrium, we divide all SNPs into blocks of length 20. Within each block, we first sample *x*_1_ ~ Bernoulli(0.5). Then we form a single haplotype block of length 20 by the following Markov chain procedure:

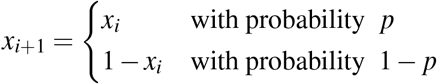

with default *p* = 0.75. For each block we form a pool of 20 haplotypes using this procedure, ensuring every one of the 40 alleles (2 at each SNP) are represented at least once. For each person, the genotype vector in a block is formed by sampling 2 haplotypes with replacement from the pool and summing the number of minor alleles at each SNP.

Depending on the simulation, the number of subjects range from 1,000 to 120,000, and the number of independent SNPs range from 10,000 to 1,000,000. We simulate data under four GLM distributions: normal (Gaussian), Bernoulli, Poisson, and negative binomial. We generate component *y_i_* of the response vector **y** by sampling from the corresponding distribution with mean 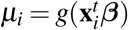, where *g* is the inverse link function. For normal models we assume unit variance, and for negative binomial models we assume 10 required failures. To avoid overflows, we clamp the mean 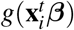 to stay within [−20, 20]. (See Ad Hoc Tactics for a detailed explanation). We apply the canonical link for each distribution, except for the negative binomial, where we apply the log link.

### 6.2 Real Data’s Quality Control Procedures

#### UK Biobank

Following the UK biobank’s own quality control procedures, we first filtered all samples for sex discordance and high heterozygosity/missingness. Second, we included only people of European ancestry and excluded first and second-degree relatives based on empiric kinship coefficients. Third, we also excluded people who had taken hypertension related medications at baseline. Finally, we only included people with ≥ 98% genotyping success rate over all chromosomes and SNPs with ≥ 99% genotyping success rate. Calculation of kinship coefficients and filtering were carried out via the OpenMendel modules `SnpArrays` (52). Remaining missing genotypes were imputed using modal genotypes at each SNP. After these quality control procedures, our UK biobank data is the same data that was used in (18).

#### Northern Finland Birth Cohort

We imputed missing genotypes with Mendel (24). Following (22), we excluded subjects with missing phenotypes, fasting subjects, and subjects on diabetes medication. We conducted quality control measures using the OpenMendel module `SnpArrays` (52). Based on these measures, we excluded SNPs with minor allele frequency ≤ 0.01 and Hardy Weinberg equilibrium p-values ≤ 10^−5^. As for non-genetic predictors, we included sex (the `sexOCPG` factor defined in (36)) as well as the first 2 principal components of the genotype matrix computed via `PLINK 2.0 alpha` (11). To put predictors, genetic and non-genetic, on an equal footing, we standardized all predictors to have mean zero and unit variance.

### 6.3 Linear Algebra with Compressed Genotype Files

The genotype count matrix stores minor allele counts. The PLINK genotype compression protocol (11) compactly stores the corresponding 0’s, 1’s, and 2’s in 2 bits per SNP, achieving a compression ratio of 32:1 compared to storage as floating point numbers. For a sparsity level *k* model, we use OpenBLAS (a highly optimized linear algebra library) to compute predicted values. This requires transforming the *k* pertinent columns of **X** into a floating point matrix **X**_*k*_ and multiplying it times the corresponding entries ***β***_*k*_ of ***β***. The inverse link is then applied to **X**_*k*_***β***_*k*_ to give the mean vector ***μ*** = *g*(**X**_*k*_***β***_*k*_). In computing the GLM gradient (equation 3.3), formation of the vector **W**_1_(**y** – ***μ***) involves no matrix multiplications. Computation of the gradient **X**^*t*^ **W**_1_(**y** – ***μ***) is more complicated because the full matrix **X** can no longer be avoided. Fortunately, the OpenMendel module `SnpArrays` (52) can be invoked to perform compressed matrix times vector multiplication. Calculation of the steplength of IHT requires computation of the quadratic form ∇*L*(***β***_*n*_)^*t*^**X**^*t*^**W**_2_**X**∇*L*(***β***_*n*_). Given the gradient, this computation requires a single compressed matrix times vector multiplication. Finally, good statistical practice calls for standardizing covariates. To standardize the genotype counts for SNP *j*, we estimate its minor allele frequency *p_j_* and then substitute the ratio 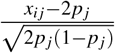 for the genotype count *x_ij_* for person *i* at SNP *j*. This procedure is predicated on a binomial distribution for the count *x_ij_*. Our previous paper (22) shows how to accommodate standardization in the matrix operations of IHT without actually forming or storing the standardized matrix.

Although multiplication via the OpenMendel module `SnpArrays` (52) is slower than OpenBLAS multiplication on small data sets, it can be as much as 10 times faster on large data sets. OpenBLAS has advantages in parallelization, but it requires floating point arrays. Once the genotype matrix **X** exceeds the memory available in RAM, expensive data swapping between RAM and hard disk memory sets in. This dramatically slows matrix multiplication. `SnpArrays` is less vulnerable to this hazard owing to compression. Once compressed data exceeds RAM, `SnpArrays` also succumbs to the swapping problem. Current laptop and desktop computers seldom have more than 32 GB of RAM, so we must resort to cluster or cloud computing when input files exceed 32 GB.

### 6.4 Computations Involving Non-genetic Covariates

Non-genetic covariates are stored as double or single precision floating point entries in an *n* × *r* design matrix **Z**. To accommodate an intercept, the first column should be a vector of 1’s. Let *γ* denote the *r* vector of regression coefficients corresponding to **Z**. The full design matrix is the block matrix (**XZ**). Matrix multiplications involving (**XZ**) should be carried out via

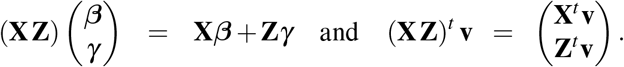

Adherence to these rules ensures a low memory footprint. Multiplication involving **X** can be conducted as previously explained. Multiplication involving **Z** can revert to BLAS.

### 6.5 Parallel Computation

The OpenBLAS library accessed by Julia is inherently parallel. Beyond that we incorporate parallel processing in cross validation. Recall that in *q*-fold cross validation we separate subjects into *q* disjoint subsets. We then fit a training model using *q* − 1 of those subsets on all desired sparsity levels and record the mean-squared prediction error on the omitted subset. Each of the *q* subsets serve as the testing set exactly once. Testing error is averaged across the different folds for each sparsity levels *k*. The lowest average testing error determines the recommended sparsity.

`MendelIHT.jl` offers 2 parallelism strategies in cross validation. Either the *q* training sets are each loaded to *q* different CPUs where each compute and test differ sparsity levels sequentially, or each of the *q* training sets are cycled through sequentially and each sparsity parameter is fitted and tested in parallel. The former tactic requires enough disk space and RAM to store *q* different training data (where each typically require (*q* − 1)/*q* GB of the full data), but offers immense parallel power because one can assign different computers to handle different sparsity levels. This tactic allows one to fit biobank scale data in less than a day assuming enough storage space and computers are available. The latter tactic requires cycling through the training sets sequentially. Since intermediate data can be deleted, the tactic only requires enough disk space and RAM to store 1 copy of the training set. `MendelIHT.jl` uses one of Julia’s (6) standard library `Distributed.jl` to achieve the aforementioned parallel strategies.

### 6.6 Ad Hoc Tactics to Prevent Overflows

In Poisson and negative binomial regressions, the inverse link argument 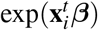 experiences numerical overflows when the inner product 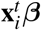 is too large. In general, we avoid running Poisson regression when response means are large. In this regime a normal approximation is preferred. As a safety feature, `MendelIHT.jl` clamps values of 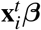 to the interval [−20, 20]. Note that penalized regression suffers from the same overflow catastrophes.

### 6.7 Convergence and Backtracking

For each proposed IHT step we check whether the objective *L*(***β***) increases. When it does not, we step-halve at most 5 times to restore the ascent property. Convergence is declared when

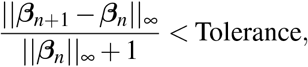

with the default tolerance being 0.0001. The addition of 1 in the denominator of the convergence criterion guards against division by 0.

## 7 Availability of source code

**Project name**: MendelIHT

**Project home page**: https://github.com/OpenMendel/MendelIHT.jl

**Operating systems**: Mac OS, Linux, Windows

**Programming language**: Julia 1.0, 1.2

**License**: MIT

The code to generate simulated data, as well as their subsequent analysis, are available in our github repository under *figures* folder. Project.toml and Manifest.toml files can be used together to instantiate the same computing environment in our paper. Notably, `MendelIHT.jl` interfaces with the OpenMendel (54) package `SnpArrays.jl` (52) and JuliaStats’s packages `Distribution.jl` (5) and `GLM.jl` (25).

## 8 Availability of supporting data and materials

The Northern Finland Birth Cohort 1966 (NFBC1966) (36) was downloaded from dbGaP under dataset accession pht002005.v1.p1. UK Biobank data are retrieved under Project ID: 48152 and 15678.

## 9 Declarations

### 9.1 List of abbreviations

GWAS: genome wide association studies;
SNP: single nucleotide polymorphism;
IHT: iterative hard threhsolding;
GLM: generalized linear models;
LD: linkage disequilibrium;
MAF: minor allele frequency;
Neg Bin: negative binomial;
NFBC: northern finland birth cohort;
HDL: high density lipoprotein;
LDL: low density lipoprotein;
SBP: systolic blood pressure;
DBP: diastolic blood pressure;

### 9.2 Ethics, Consent for publication, competing interest

The authors declare no conflicts of interest. As described in (36), informed consent from all study subjects of NFBC1966 was obtained using protocols approved by the Ethical Committee of the Northern Ostrobothnia Hospital District.

### 9.3 Funding

Benjamin Chu was supported by NIH T32-HG002536 training grant and the 2018 Google Summer of Code. Kevin Keys was supported by a diversity supplement to NHLBI grant R01HL135156, the UCSF Bakar Computational Health Sciences Institute, the Gordon and Betty More Foundation grant GBMF3834, and the Alfred P. Sloan Foundation grant 2013-10-27 to UC Berkeley through the Moore-Sloan Data Sciences Environment initiative at the Berkeley Institute for Data Science (BIDS). Kenneth Lange and Hua Zhou were supported by grants from the National Human Genome Research Institute (HG006139) and the National Institute of General Medical Sciences (GM053275). Janet Sinsheimer was supported by grants from the National Institute of General Medical Sciences (GM053275), the National Human Genome Research Institute (HG009120) and the National Science Foundation (DMS-1264153). Christopher German was supported by the Burroughs Wellcome Fund Inter-school Training Program in Chronic Diseases (BWF-CHIP).

### 9.4 Author’s Contributions

JSS, KK, KL, ES, BC contributed to the design of the study, interpretation of results, and writing of the original draft manuscript. BC designed and implemented the simulations and conducted the data analyses. HZ, CG, and JZ contributed to the analysis of UKBB results. KK and BC developed the software. KL and BC developed the algorithms. ES assisted in the comparisons to marginal GWAS. All authors have read, made suggestions and ultimately approved the final manuscript.

## 10 Acknowledgements

We thank UCLA for providing the computing resources via the Hoffman2 cluster.

The NFBC1966 Study is conducted and supported by the National Heart, Lung, and Blood Institute (NHLBI) in collaboration with the Broad Institute, UCLA, University of Oulu, and the National Institute for Health and Welfare in Finland. This manuscript was not prepared in collaboration with investigators of the NFBC1966 Study and does not necessarily reflect the opinions or views of the NFBC1966 Study Investigators, Broad Institute, UCLA, University of Oulu, National Institute for Health and Welfare in Finland and the NHLBI.

## Notes

#### Summary of Updates

+ Added Figure 4 to compare our UK Biobank analysis with a traditional GWAS method. We also compared our method to a recent publication (German et al. 2019 Genetic Epidemiology) that analyzed the same data + Added parameter estimation comparisons of our method with traditional GWAS in Table 3.

https://github.com/OpenMendel/MendelIHT.jl

## References

[1] Gad Abraham, Yixuan Qiu, and Michael Inouye. FlashPCA: principal component analysis of Biobank-scale genotype datasets. Bioinformatics, 2017.

[2] David H. Alexander and Kenneth Lange. Stability selection for genome-wide association. Genetic Epidemiology, 35:722–728, 2011.

[3] Amir Beck. Introduction to nonlinear optimization: Theory, algorithms, and applications with MATLAB, volume 19. Siam, 2014.

[4] Amir Beck and Marc Teboulle. A linearly convergent algorithm for solving a class of nonconvex/affine feasibility problems. In Fixed-Point Algorithms for Inverse Problems in Science and Engineering, pages 33–48. Springer, 2011.

[5] Mathieu Besançon, David Anthoff, Alex Arslan, Simon Byrne, Dahua Lin, Theodore Papamarkou, and John Pearson. Distributions.jl: Definition and modeling of probability distributions in the juliastats ecosystem. arXiv e-prints, page arXiv:1907.08611, Jul 2019.

[6] Jeff Bezanson, Alan Edelman, Stefan Karpinski, and Viral B Shah. Julia: A fresh approach to numerical computing. SIAM Review, 59:65–98, 2017.

[7] Thomas Blumensath and Mike E Davies. Normalized iterative hard thresholding: Guaranteed stability and performance. IEEE Journal of Selected Topics in Signal Processing, 4:298–309, 2010.

[8] Patrick Breheny and Jian Huang. Coordinate descent algorithms for nonconvex penalized regression, with applications to biological feature selection. Annals of Applied Statistics, 5:232–253, 2011.

[9] William S Bush and Jason H Moore. Genome-wide association studies. PLoS Computational Biology, 8:e1002822, 2012.

[10] Rita M Cantor, Kenneth Lange, and Janet S Sinsheimer. Prioritizing GWAS results: a review of statistical methods and recommendations for their application. The American Journal of Human Genetics, 86:6–22, 2010.

[11] Christopher C Chang, Carson C Chow, Laurent CAM Tellier, Shashaank Vattikuti, Shaun M Purcell, and James J Lee. Second-generation PLINK: rising to the challenge of larger and richer datasets. Giga-Science, 4:7, 2015.

[12] de Lamare, Rodrigo C. Knowledge-aided normalized iterative hard thresholding algorithms and applications to sparse reconstruction. arXiv preprint arXiv:1809.09281, 2018.

[13] Annette J Dobson and Adrian Barnett. An introduction to generalized linear models. Chapman and Hall/CRC, 2008.

[14] Simon Foucart. Hard thresholding pursuit: an algorithm for compressive sensing. SIAM Journal on Numerical Analysis, 49:2543–2563, 2011.

[15] Jerome Friedman, Trevor Hastie, and Rob Tibshirani. Regularization paths for generalized linear models via coordinate descent. Journal of Statistical Software, 33:1, 2010.

[16] Jerome Friedman, Trevor Hastie, and Robert Tibshirani. A note on the group lasso and a sparse group lasso. arXiv preprint arXiv:1001.0736, 2010.

[17] Lisa Gai and Eleazar Eskin. Finding associated variants in genome-wide association studies on multiple traits. Bioinformatics, 34:i467–i474, 2018.

[18] Christopher A German, Janet S Sinsheimer, Yann C Klimentidis, Hua Zhou, and Jin J Zhou. Ordered multinomial regression for genetic association analysis of ordinal phenotypes at Biobank scale. Genetic Epidemiology, 2019.

[19] Christopher A German and Hua Zhou. MendelPlots.jl: Julia package for plotting results from GWAS, February 2020.

[20] Buhm Han and Eleazar Eskin. Random-effects model aimed at discovering associations in meta-analysis of genome-wide association studies. The American Journal of Human Genetics, 88:586–598, 2011.

[21] Gabriel E. Hoffman, Benjamin A. Logsdon, and Jason G. Mezey. PUMA: A unified framework for penalized multiple regression analysis of GWAS data. PLoS Computational Biology, 9:e1003101, 06 2013.

[22] Kevin L Keys, Gary K Chen, and Kenneth Lange. Iterative hard thresholding for model selection in genome-wide association studies. Genetic Epidemiology, 41:756–768, 2017.

[23] Kenneth Lange. Numerical analysis for statisticians. Springer Science & Business Media, 2010.

[24] Kenneth Lange, Jeanette C Papp, Janet S Sinsheimer, Ram Sripracha, Hua Zhou, and Eric M Sobel. Mendel: the swiss army knife of genetic analysis programs. Bioinformatics, 29:1568–1570, 2013.

[25] Dahua Lin, John Myles White, Simon Byrne, Douglas Bates, Andreas Noack, John Pearson, Alex Arslan, Kevin Squire, David Anthoff, Theodore Papamarkou, Mathieu Besançon, and et al. JuliaStats/Distributions.jl: a Julia package for probability distributions and associated functions, may 2019.

[26] Po-Ru Loh, George Tucker, Brendan K Bulik-Sullivan, Bjarni J Vilhjalmsson, Hilary K Finucane, Rany M Salem, Daniel I Chasman, Paul M Ridker, Benjamin M Neale, Bonnie Berger, Nick Patterson, and Alkes L Price. Efficient bayesian mixed-model analysis increases association power in large cohorts. Nature Genetics, 47:284, 2015.

[27] Jacqueline MacArthur, Emily Bowler, Maria Cerezo, Laurent Gil, Peggy Hall, Emma Hastings, Heather Junkins, Aoife McMahon, Annalisa Milano, Joannella Morales, and et al. The new NHGRI-EBI Catalog of published genome-wide association studies (GWAS Catalog). Nucleic acids research, 45:D896–D901, 2016.

[28] Alicia R Martin, Konrad J Karczewski, Sini Kerminen, Mitja I Kurki, Antti-Pekka Sarin, Mykyta Artomov, Johan G Eriksson, Tõnu Esko, Giulio Genovese, Aki S Havulinna, and et al. Haplotype sharing provides insights into fine-scale population history and disease in finland. The American Journal of Human Genetics, 102:760–775, 2018.

[29] Rahul Mazumder, Jerome H. Friedman, and Trevor Hastie. SparseNet: Coordinate descent with non-convex penalties. Journal of the American Statistical Association, 106:1125–1138, 2011.

[30] Peter McCullagh. Generalized Linear Models. Routledge, 2018.

[31] Lukas Meier, Sara Van De Geer, and Peter Bühlmann. The group lasso for logistic regression. Journal of the Royal Statistical Society: Series B (Statistical Methodology), 70:53–71, 2008.

[32] Nicolai Meinshausen and Peter Bühlmann. Stability selection. Journal of the Royal Statistical Society: Series B (Statistical Methodology), 72:417–473, 2010.

[33] Stacey Melquist, David W Craig, Matthew J Huentelman, Richard Crook, John V Pearson, Matt Baker, Victoria L Zismann, Jennifer Gass, Jennifer Adamson, Szabolcs Szelinger, et al. Identification of a novel risk locus for progressive supranuclear palsy by a pooled genomewide scan of 500,288 single-nucleotide polymorphisms. The American Journal of Human Genetics, 80:769–778, 2007.

[34] Deanna Needell and Joel A Tropp. Cosamp: Iterative signal recovery from incomplete and inaccurate samples. Applied and Computational Harmonic Analysis, 26:301–321, 2009.

[35] SMA Khaleelur Rahman, M Mohamed Sathik, and K Senthamarai Kannan. Multiple linear regression models in outlier detection. International Journal of Research in Computer Science, 2(2):23, 2012.

[36] Chiara Sabatti, Susan K Service, Anna-Liisa Hartikainen, Anneli Pouta, Samuli Ripatti, Jae Brodsky, Chris G Jones, Noah A Zaitlen, Teppo Varilo, Marika Kaakinen, and et al. Genome-wide association analysis of metabolic traits in a birth cohort from a founder population. Nature genetics, 41:35, 2009.

[37] Armin P Schoech, Daniel M Jordan, Po-Ru Loh, Steven Gazal, Luke J O’Connor, Daniel J Balick, Pier F Palamara, Hilary K Finucane, Shamil R Sunyaev, and Alkes L Price. Quantification of frequencydependent genetic architectures in 25 uk biobank traits reveals action of negative selection. Nature communications, 10(1):790, 2019.

[38] Cathie Sudlow, John Gallacher, Naomi Allen, Valerie Beral, Paul Burton, John Danesh, Paul Downey, Paul Elliott, Jane Green, Martin Landray, Bette Liu, Paul Matthews, Giok Ong, Jill Pell, Alan Silman, Alan Young, Tim Sprosen, Tim Peakman, and Rory Collins. Uk biobank: an open access resource for identifying the causes of a wide range of complex diseases of middle and old age. PLoS Medicine, 12:e1001779, 2015.

[39] Robert Tibshirani. Regression shrinkage and selection via the lasso. Journal of the Royal Statistical Society. Series B (Methodological), pages 267–288, 1996.

[40] Shashaank Vattikuti, James J Lee, Christopher C Chang, Stephen DH Hsu, and Carson C Chow. Applying compressed sensing to genome-wide association studies. GigaScience, 3:10, 2014.

[41] Peter M Visscher, Naomi R Wray, Qian Zhang, Pamela Sklar, Mark I McCarthy, Matthew A Brown, and Jian Yang. 10 years of GWAS discovery: biology, function, and translation. The American Journal of Human Genetics, 101:5–22, 2017.

[42] Paul K Whelton, Robert M Carey, Wilbert S Aronow, Donald E Casey, Karen J Collins, Cheryl Dennison Himmelfarb, Sondra M DePalma, Samuel Gidding, Kenneth A Jamerson, Daniel W Jones, et al. 2017 ACC/AHA/AAPA/ABC/ACPM/AGS/APhA/ASH/ASPC/NMA/PCNA guideline for the prevention, detection, evaluation, and management of high blood pressure in adults: a report of the American College of Cardiology/American Heart Association Task Force on Clinical Practice Guidelines. Journal of the American College of Cardiology, 71(19):e127–e248, 2018.

[43] Tong Tong Wu, Yi Fang Chen, Trevor Hastie, Eric Sobel, and Kenneth Lange. Genome-wide association analysis by lasso penalized logistic regression. Bioinformatics, 25:714–721, 2009.

[44] Tong Tong Wu and Kenneth Lange. Coordinate descent algorithms for lasso penalized regression. The Annals of Applied Statistics, 2:224–244, 2008.

[45] Jason Xu, Eric Chi, and Kenneth Lange. Generalized Linear Model Regression under Distance-to-set Penalties. In Advances in Neural Information Processing Systems 30., pages 1385–1395. Curran Associates, Inc., 2017.

[46] Fan Yang, Rina Foygel Barber, Prateek Jain, and John Lafferty. Selective inference for group-sparse linear models. In Advances in Neural Information Processing Systems, pages 2469–2477, 2016.

[47] Xiao-Tong Yuan, Ping Li, and Tong Zhang. Gradient hard thresholding pursuit. Journal of Machine Learning Research, 18:166–1, 2017.

[48] Achim Zeileis, Christian Kleiber, and Simon Jackman. Regression models for count data in R. Journal of Statistical Software, 27:1–25, 2008.

[49] Jian Zeng, Ronald De Vlaming, Yang Wu, Matthew R Robinson, Luke R Lloyd-Jones, Loic Yengo, Chloe X Yap, Angli Xue, Julia Sidorenko, Allan F McRae, et al. Signatures of negative selection in the genetic architecture of human complex traits. Nature genetics, 50(5):746, 2018.

[50] Tong Zhang. Analysis of multi-stage convex relaxation for sparse regularization. Journal of Machine Learning Research, 11:1081–1107, 2010.

[51] Hua Zhou, David H Alexander, Mary E Sehl, Janet S Sinsheimer, Eric M Sobel, and Kenneth Lange. Penalized regression for genome-wide association screening of sequence data. In Pacific Symposium on Biocomputing, pages 106–117. World Scientific, 2011.

[52] Hua Zhou, Jeanette C Papp, Seyoon Ko, Christopher A German, Joshua Day, Marc A Suchard, Alfonso Landeros, and Andreas Noack. SnpArrays.jl: Julia package for compressed storage of SNP data, February 2020.

[53] Hua Zhou, Mary E Sehl, Janet S Sinsheimer, and Kenneth Lange. Association screening of common and rare genetic variants by penalized regression. Bioinformatics, 26:2375, 2010.

[54] Hua Zhou, Janet S Sinsheimer, Douglas M Bates, Benjamin B Chu, Christopher A German, Sarah S Ji, Kevin L Keys, Juhyun Kim, Seyoon Ko, Gordon D Mosher, and et al. OpenMendel: a cooperative programming project for statistical genetics. Human Genetics, pages 1–11, 2019.

